# Oyster aquaculture enhances sediment microbial diversity– Insights from a multi-omics study

**DOI:** 10.1101/2023.11.13.566866

**Authors:** Joshua T.E. Stevens, Nicholas E. Ray, Alia N. Al-Haj, Robinson W. Fulweiler, Priyanka Roy Chowdhury

## Abstract

The global aquaculture industry has grown substantially, with consequences for coastal ecology and biogeochemistry. Oyster aquaculture can alter the availability of resources for microbes that live in sediments as oysters move large quantities of organic material to the sediments via filter feeding, possibly leading to changes in the structure and function of sediment microbial communities. Here, we use a chronosequence approach to investigate the impacts of oyster farming on sediment microbial communities over 7 years of aquaculture activity in a temperate coastal system. We detected shifts in bacterial composition (16S rRNA amplicon sequencing), changes in gene expression (meta-transcriptomics), and variations in sediment elemental concentrations (sediment geochemistry) across different durations of oyster farming. Our results indicate that both the structure and function of bacterial communities vary between control (no oysters) and farm sites, with an overall increase in diversity and a shift towards anoxic tolerance in farm sites. However, little to no variation was observed in either structure or function with respect to farming duration suggesting these sediment microbial communities are resilient to change. We also did not find any significant impact of farming on heavy metal accumulation in the sediments. The minimal influence of long-term oyster farming on sediment bacterial function and biogeochemical processes as observed here can bear important consequences for establishing best practices for sustainable farming in these areas.

**Importance:** Sediment microbial communities drive a range of important ecosystem processes such as nutrient recycling and filtration. Oysters are well-known ecological engineers, and their presence is increasing as aquaculture expands in coastal waters globally. Determining how oyster aquaculture impacts sediment microbial processes is key to understanding current and future estuarine biogeochemical processes. Here, we use a multi-omics approach to study the effect of different durations of oyster farming on the structure and function of bacteria and elemental accumulation in the farm sediments. Our results indicate an increase in the diversity of bacterial communities in the farm sites with no such increases observed for elemental concentrations. Further, these effects persist across multiple years of farming with an increase of anoxic tolerant bacteria at farm sites. The multi-omics approach used in this study can serve as a valuable tool to facilitate understanding of the environmental impacts of oyster aquaculture.

## Introduction

Oysters are filter-feeders that regulate biogeochemical processes in coastal ecosystems by delivering organic material (OM) to sediments and excreting dissolved nutrients. Once common in many coastal systems around the world, now over harvest and diseases have made natural oyster populations only a small fraction of their former historic extent (1–4). Today, oyster aquaculture is expanding rapidly, particularly on the east coast of the United States (5–6). In addition to the economic value of oyster aquaculture, oysters are ecosystem engineers, providing a host of benefits including habitat provisioning and storm surge protection (7). Their high filtration capacity (∼ 5 L of water per hour) enables them to filter large amounts of particulate matter out of the water. These particles have two fates – they can be ingested and deposited to the sediments below as feces, or rejected, wrapped in mucus, and deposited as pseudofeces. The deposition of this OM stimulates a range of microbial processes by changing the availability of compounds used in microbial metabolism (8), altering oxygen penetration into sediments (9), and possibly promoting the build-up of chemicals that inhibit certain metabolic pathways (8, 10). Further, in a simulated lab study Newell et al. (11) suggested that most aquaculture driven changes in nutrient fluxes presumably occur through intermediate microbial processes. Taken together, these studies indicate that the response of sediment microbial communities to the pressure of oyster-mediated OM loading will have important implications for biogeochemistry in the system.

To date, most of the research on the effect of oysters on coastal geochemistry have focused on sediment nitrogen (N) cycling, specifically addressing how oysters alter rates of individual N cycling pathways and the net exchange of various N compounds between the sediment and water column (12). Underlying these changes and individual processes are various microbial guilds that may change in abundance or activity in response to the presence of oysters. For example, oyster farms are considered ‘hot spots’ for N removal via microbial denitrification (13, 14), the microbial conversion of biologically active nitrogen to unreactive di-nitrogen gas. Microbial denitrification rates are influenced by a variety of factors including the quality and quantity of organic matter (15, 16), oxygen concentrations (17), and nitrate availability (18) – all factors impacted by oyster farming (10, 19). Changes in N cycling can also initiate or coincide with changes in the availability and concentration of carbon (C), phosphorus (P), and sulfur (S) (20). In areas of high OM deposition, such as beneath oyster cages, hydrogen sulfide (H_2_S) can build up, subsequently inhibiting rates of nitrification and denitrification (21, 22). In addition to these 4 elements (C, N, P and S), oyster mediated OM loading can also enhance the availability of heavy (such as Cu, Zn, and Pb: 23-25) and trace metals (such as As, Cd, Co, Cr, Fe, Mn, and Ni: 26, 27), several of which are required in enzymes that perform microbial metabolism. For example, enzymes important for denitrification such as *nitrate reductase,* requires a molybdenum (Mo) protein cofactor for reducing nitrate to nitrite (28) and *nitrous oxide reductase* requires copper (Cu) for reducing nitrous oxide (N_2_O) to di-nitrogen (N_2_) (29). The concentration of iron (Fe) has a significant role in mediating the microbial mobilization of iron-bound phosphorus in marine sediment (30). Therefore, it can be assumed that any changes in the environmental supply of these metals (e.g., Mo, Cu and Fe) may significantly impact the cycling of other nutrients (e.g., N and P) (31). Considering the effect of oyster farming on both the availability of metals and the microbial communities is an important next step toward broader insights into how oysters regulate biogeochemical processes across systems.

Past studies have investigated how sediment microbial communities may differ between oyster habitats and nearby areas without oysters. For example, Feinman et. al., (32) demonstrated higher bacterial abundance beneath oyster aquaculture compared to bare sediment and there is also evidence for significant shifts in communities following implementation of other types of shellfish culturing (21, 33, 34). While these studies provide insight into how sediment communities may change following introduction of shellfish aquaculture, they have focused on the dynamics of a few well characterized microbial groups, and do not capture the complexities in communal shifts in response to farming durations. Nor do they capture the impact of related elemental biogeochemical processes that are important for controlling the microbial driven nutrient cycles in sediments.

Bacteria can respond promptly to changes in their environment due to shorter generation time allowing faster rates of evolutionary changes. For example, bacterial communities in restored salt marshes have been found to resemble those of reference salt marshes before there was evidence of plant growth (35), making bacterial communities’ effective ecological indicators. Given the potential for rapid microbial evolution, community composition alone may not accurately capture all functional variations that arises in response to environmental impacts. Therefore, an integrative approach comparing community wide changes in structure and function of bacteria is vital for a holistic understanding of the extent of farming impacts, which in turn can be useful in future monitoring and management of these farms. In this study, we use a multi-omics approach to test shifts in sediment bacterial communities along an oyster aquaculture ‘chronosequence’ (space-for-time substitution; 36). To that end, we first used 16S rRNA amplicon sequencing to characterize changes in the structure and function of bacterial communities in bare sediments and sediments beneath oyster farms of various ages. Next, we used RNA meta-transcriptomics to identify similar shifts in response to oyster farming. Using both DNA and RNA sequencing allowed us to detect changes in the “potential” (DNA) compared to the “active” (RNA) shifts in response to farming. Finally, we measured changes in sediment elemental concentrations (i.e., ionome; 37) in relation to the age of oyster farming to identify the effects of farming on elements beyond N, C, P and S. We hypothesized that there would be significant changes in both the structure and function of the bacterial community in response to farming duration due to changes in both biotic and abiotic factors induced by the oysters. Functional data allowed us to identify specific bacterial processes and their associated functions that mediates the above shifts in composition. Based on previous studies (38, 39), we predict greater accumulation of metals (e.g., Cd, Cu, Pb, Zn) in farmed sites due to OM deposition by the oysters. We also predict a differential effect of elements (owing to variations in concentrations) on bacterial communities in control versus farmed sites.

## Materials and Methods

### Study Site and Sample Collection

Sediments were collected from Ninigret Pond (41.357°N, 71.6534°E), a coastal back barrier lagoon located in southern Rhode Island, USA (Appendix S1). 4 sites were selected within a commercial oyster farm (0.016 km^2^) in the lagoon with varying durations of oyster farming (0, 3, 5, and 7 years) that corresponded to a chrono-sequence (space-for-time substitution). In June of 2015, duplicate sediment samples were collected using a 25-mm corer from the top 1 cm of sediments under oyster cages from 3 locations within each of the 4 sites, yielding 24 total samples. Sediment samples were placed on dry ice to be transported back to the laboratory and stored at −80 °C until downstream analyses. A more detailed description of the study location, oyster farming approach, and methods for sample collection can be found in Ray et al. (36).

### DNA extraction and analyses

We used 16S rRNA amplicon sequencing to identify changes in bacterial community composition based on the protocol described in Stevens et. al. (40). Briefly, DNA were extracted using the Qiagen DNeasy PowerSoil pro kit (Cat. No. 47014). Purified DNA was then shipped to MR DNA (Shallowater, TX; https://www.mrdnalab.com/) for sequencing on an Ion Torrent personal genome machine (PGM) after PCR amplification of the 16S rRNA V4 region with primers 515F and 806R (41). Initial quality control of sequences was done using the MR DNA analysis pipeline to remove primers, barcodes, and any sequences <150 bp. A 97% similarity cutoff was applied to filtered reads to identify Operational taxonomic units (OTUs) and chimeras were removed using a modified UCHIME algorithm (42). OTUs were clustered using USEARCH and classified using BLASTN against databases derived from the Ribosomal Database Project II (*v 11.5*; http://rdp.cme.msu.edu; 43) and NCBI database (https://www.ncbi.nlm.nih.gov).

### RNA extraction and analyses

We isolated RNA from 6 sediment samples (3 replicates each from the control and 7-year old site). RNA from all 6 samples were extracted from ∼2 gm of sediment using Qiagen RNeasy PowerSoil Total RNA Kit (Cat. No. 12866) following the manufacturer’s protocol. RNA was quantified using the Qubit RNA HR assay kit (Thermo Fisher Scientific Inc., Waltham, MA, USA) in a Qubit Flurometer (Thermo Fisher, Cambridge, MA), before shipping to UNH Hubbard Center for Genome studies for sequencing (https://hcgs.unh.edu/). RNA was reverse transcribed using the SuperScript Double-Stranded cDNA Synthesis Kit (Thermo Fisher, Cambridge, MA). Illumina compatible Nextera DNA Flex Library Prep Kit (Cat. No. 20018705) was used to construct the RNA libraries following the manufacturer’s protocol. Illumina-compatible adapters from the Nextera DNA Unique Dual Indexes (Cat. No. 20027213) were used to attach individual bar codes to all 6 libraries. Library size distribution was determined using a Bioanalyzer 2100 (Agilent Technologies, Santa Clara CA, USA) with DNA High-Sensitivity chips and reagents (Agilent Technologies, Santa Clara CA, USA). Illumina TruSeq SBS v4 reagent kit (300 cycles) was used to generate paired-end 150 bp reads using the Illumina HiSeq platform (Illumina, San Diego CA, USA). Initial quality check was done with FastQC (v0.11.6) (<u>bioinformatics.babraham.ac.uk/projects/fastqc</u>). Low quality reads and TruSeq3-PE adapters were then removed in Trimmomatic (v 0.27) using default settings (44). Trimmed and filtered reads were uploaded to the MG-RAST API server (http://metagenomics.anl.gov) for both taxonomic and functional analyses of MetaT data following previously established protocol (45, 46). Briefly, RNA was first converted to cDNA. A minimum e-value <1e^−5^ and 60% identity cut-off were used for database searches within MG-RAST. RefSeq database was used to tabulate sequence counts at each taxonomic levels and functional genes from the cDNA library were annotated with SEED Subsystem and visualized using KEGG (an internal tool based on the Kyoto Encyclopedia of Genes and Genomes pathway mapping system).

### Metal Extraction and Analyses

From each of the 24 sediment samples, thawed sediments were grounded to a fine powder using a mortar and pestle. ∼2 g of each dried and thawed sample was then sent to the Trace Metal Analyses Core in Dartmouth College to measure elemental concentrations using an Agilent™ 7900 inductively coupled plasma mass spectrometer (ICP-MS). Prior to analysis, samples were digested in 5 mL HNO_3_: HCL (9:1) mixture and heated at 110 °C for an hour. On cooling, 45 mL of DI water was added before running in the ICP-MS with a commercial internal standard mix for calibration purposes.

### Statistical data analyses

All analyses were performed in the statistical platform R using the RStudio interface (*v 4.2.1*) (R Core Team, 2017), including *vegan* (47), *ggplot2* (48) and *DESeq2* packages (49). The *rarefy* function in the *vegan* package was used to level unequal sequencing depths across amplicon samples. Alpha diversity (Shannon Index) was evaluated using the *vegan* package and Kruskal-Wallis non-parametric test (pairwise) was used to test differences between sites. Beta diversity (sample clustering) was visualized by non-metric multidimensional scaling (nMDS) ordination using Bray-Curtis dissimilarity matrix in *vegan* (Final Stress: 0.123). Significance of microbial community shifts within and between samples were tested by ANOSIM and PERMANOVA respectively, using the *adonis ()* function in the *vegan* package. Linear discriminant analysis (LDA) effect size (LEfSe) analysis was used to detect site-specific bacterial markers using LEfSe of Galaxy of the Huttenhower lab (50). Potential bacterial functions were identified from the 16S data using PICRUSt (51) in Qiime2 (52). Differential abundance in KEGG functional pathways were determined in *ggpicrust2* (53) using ALDEx2 (54) between control and farmed sites that uses Wilcoxon rank test for statistical significance between treatments. All predictions were corrected for multiple testing (Benjamini–Hochberg method, FDR q < 0.05).

For sediment elemental concentrations, nMDS based on Bray-Curtis dissimilarity matrix (Final Stress: 0.163) were performed on z-transformed data to normalize distributions across samples, followed by PERMANOVA and ANOSIM tests for differences among sites. Pearson’s correlation analysis using the *stats* package was used to assess associations between elements. Individual concentrations of elements were compared across sites using a one-way ANOVA followed by Tukeys post-hoc tests. The contribution of elements on the variations in bacterial community was estimated using distance-based redundancy analyses (db-RDA) using *rda()* function in *vegan*. Only elements that showed a significant correlation with grouped samples using the *envfit* function in vegan were used in the redundancy analyses (47).

For RNA sequencing, KEGG annotated gene counts and RefSeq based taxonomic counts were used for nMDS ordination using Bray-Curtis dissimilarity matrix (Final Stress: 0.006). followed by ANOSIM and PERMANOVA analyses for site-specific differences. Differential gene expression of KEGG transcripts between control and 7-year site were calculated using *DEseq2*. Results were plotted in SRplot (http://www.bioinformatics.com.cn/srplot). Diversity Indices of RNA derived taxonomic profiles were calculated using *vegan* and Kruskal-Wallis non-parametric test (pairwise) was used to test for differences between sites. To test for similarity between community composition derived from DNA and RNA sequences, a two-tailed Student’s t-test were performed on Canberra pairwise community distances at the phyla level using the *vegdist* function in *vegan* (see 55). Mantel tests were performed to identify significant associations between physical environmental parameters and OTU abundances, elemental concentrations and RNA derived taxonomic and functional gene counts. Environmental parameters were obtained from Ray et al. (56). The assumptions of normality and homogeneity of variances were met for all analyses, wherever applicable.

## Results

### DNA Community Structure & Function

16S rRNA amplicon sequencing in all 24 samples (4 sites x 6 replicates) showed a mean of 58,137 sequences per sample (±15,636 (SD); 40). We identified 13,147 OTUs belonging to 53 phyla, 119 classes, and 1137 genera across samples. Rarefaction curves (Appendix S2a) indicate that most samples had comparable sequencing depth, with all 3-year replicates showing greater OTU abundances compared to the rest. Control, 5-year and 7-year farm sediment replicates had nearly equal abundances, but the control replicates exhibited lower species richness than the farmed sites (Fig. 1A; Table 1). These findings were also supported by diversity indices analyses, where control samples were significantly less diverse than all 3 farmed sites (p < 0.001; Kruskal–Wallis test), with no considerable variation between the 3-, 5-and 7-year sites (Table 1; Fig. 1A). nMDS ordination was used to examine clustering at the OTU levels where control sites clustered separately from other farmed sites. 3-and 5-year clustered closer compared to 7-year sites (Fig. 1B).

**Figure 1:**
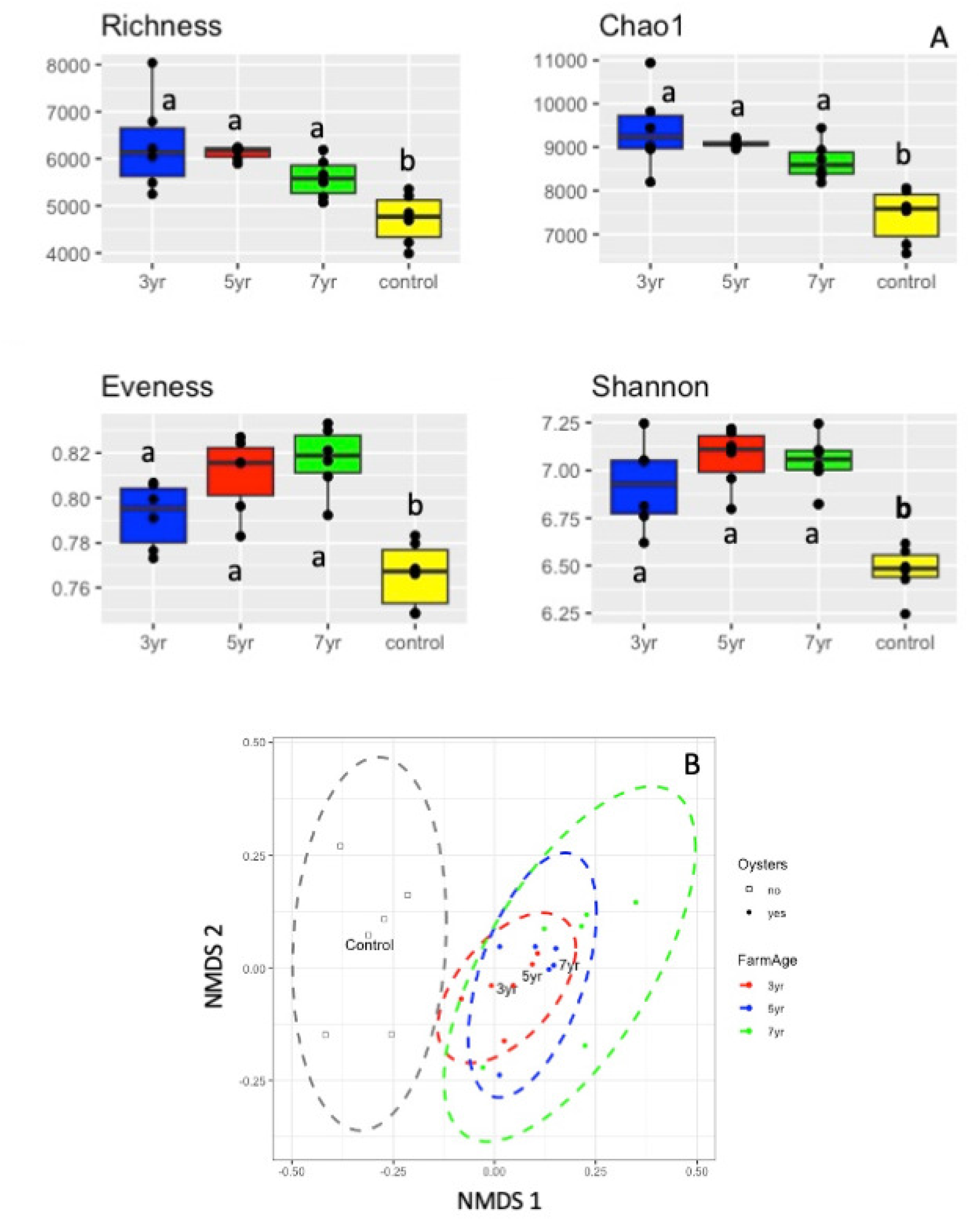
(A) Diversity Analyses and (B) Non-metric multidimensional scaling (nMDS) of 16S rRNA community based on Operational Taxonomic Unit (OTU). Different letters denote significant differences in diversity.

**Table 1:**
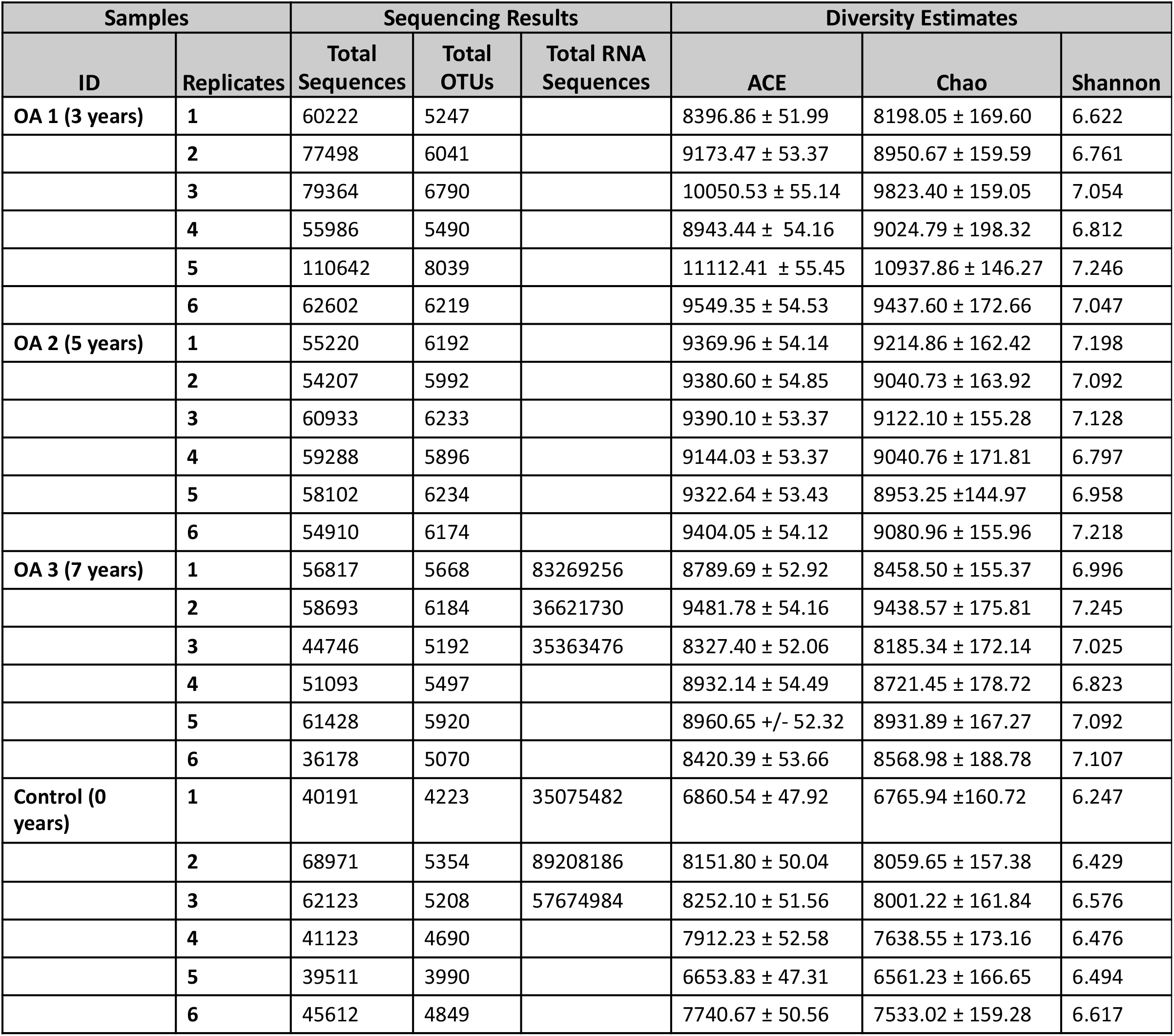
16S-rRNA gene and Illumina HiSeq Sequencing Output showing total sequences obtained in each, number of OTUs and KEGG annotations in 16s-rRNA and HiSeq mRNA respectively, and diversity indices of OTU abundances across the 24 sample sites: *ACE* abundance-based coverage estimator, *Chao* Chao’s species richness estimator, *Shannon* Shannon’s diversity index.

PERMANOVA (differences between groups) and ANOSIM (comparison of between group differences vs. within group) tests indicated a significant difference between control and farmed sites at the OTU level (Table 2 A & B). Bacterial groups that are more likely indicators of this difference were characterized by linear discriminant analysis (LDA) of effect size (LEfSe) that revealed 13 OTUs were enriched in the farmed sites (3-, 5-and 7-; LDA Scores 3.4 -5.3) and 14 OTUs were enriched in the control sites (LDA Scores 2.7 – 5.6) (Fig. 2A; Appendix S3). Taxa enriched in farm sites belonged to phylum *Proteobacteria* except one that belonged to *Firmicutes.* Within *Proteobacteria,* 11 belonged to class *Deltaproteobacteria* and 1 to *Epsilonproteobacteria*. *Desulfobacterales* and *Desulfuromonadales* were the two enriched order in the farm sites. In control sites, phyla *Proteobacteria* (8)*, Bacteriodetes* (5) *and Spirochaetes* (Class *Spirochaetia*) (1) were enriched (Appendix S3). Within P*roteobacteria*, class *Alphaproteobacteria* (order *Rhodobacterales*) showed the highest LDA score followed by *Gammaproteobacteria*. Within *Bacteriodetes*, class *Sphingobacteriia* (order *Sphingobacteriales*) had the highest LDA score followed by *Cytophagia* (order *Cytophagales*) (Fig. 2A; Appendix S3).

**Figure 2:**
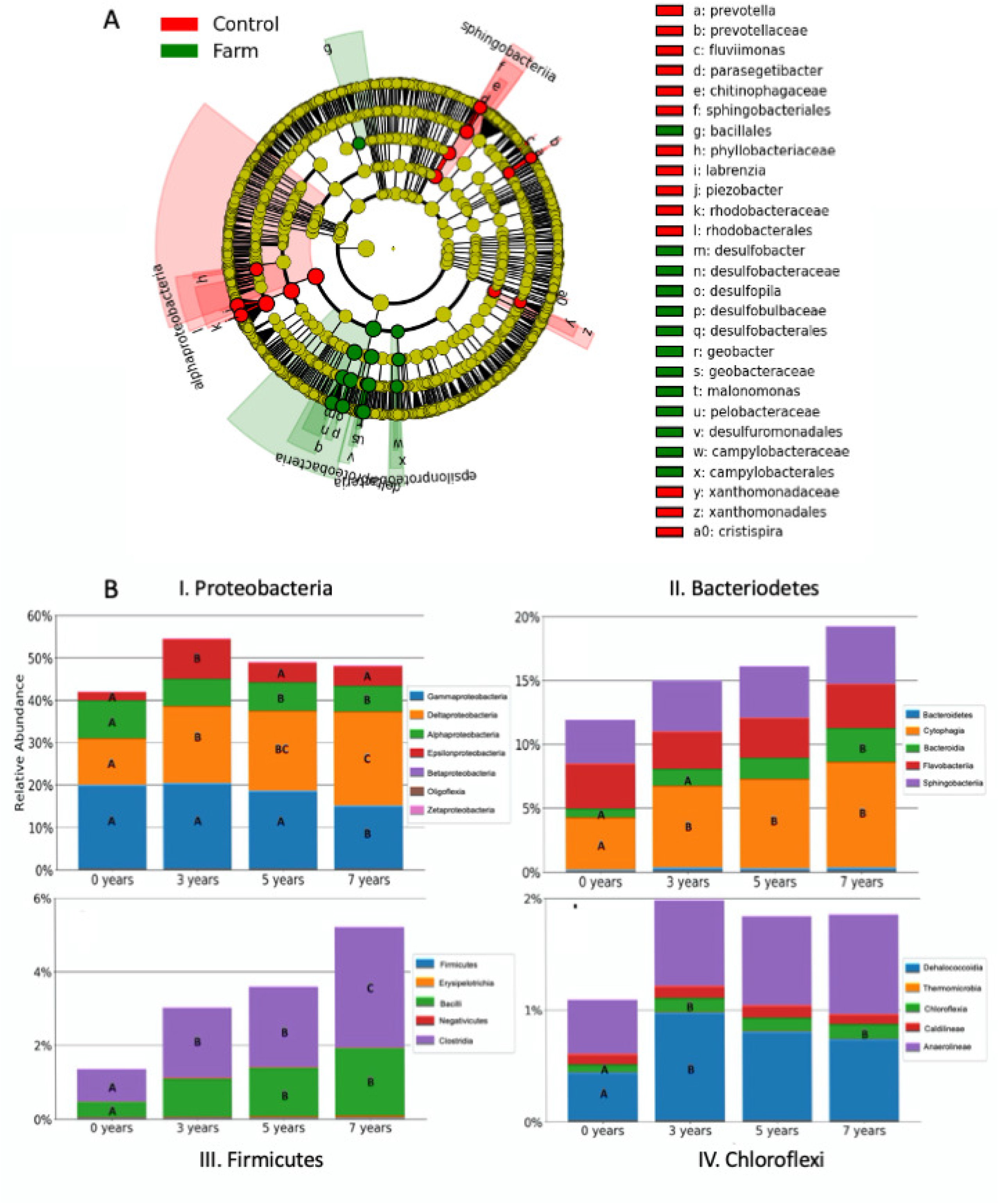
(A) Linear discriminant analysis (LDA) effect size (LEfSe) of taxonomic biomarkers in control (red) and 7-year (green) farm sites (B) Relative abundances of different classes within phylum *Proteobacteria* (I), *Bacteriodetes* (II), *Firmicutes* (III), and *Chloroflexi* (IV). Different letters across the same color bars denotes significant differential abundance.

**Table 2:** (A) Permutational multivariate analysis of variance (PERMANOVA) and (B) Analysis of Similarity (ANOSIM) table for differences in bacterial OTUs, elemental concentrations, KEGG annotated and RefSeq annotated RNA transcripts in grouped samples (control, 3-, 5-and 7-years)

| (A) PERMANOVA (Differences between Groups) |  |  |  | (B) ANOSIM (Between Group vs. Within Group) |  |  |
| --- | --- | --- | --- | --- | --- | --- |
| Source | df | F-value | p-value | Source | R-statistic | p-value |
| OTUs | 3 | 3.2063 | 0.001 *** | OTUs | 0.581 | 0.00001 *** |
| Elements | 3 | 3.3494 | 0.009 ** | Elements | 0.2994 | 0.0017 *** |
| RNA-KEGG | 2 | 0.2682 | 0.933 | RNA-KEGG | 0.3182 | 0.8833 |
| RNA-Genera | 2 | 1.3389 | 0.2167 | RNA-Genera | 0 | 0.5 |

Significance codes: 0 '\*\*\*' 0.001 '\*\*' 0.01 '\*' 0.05 '.' 0.1 ' ' 1
Number of permutations: 999

Phylum level abundances across the 4 sites are published previously in Stevens et al. (40; Genome Announcement). Several classes within the four most abundant phyla (*Proteobacteria*, *Bacteriodetes*, *Firmicutes,* and *Chloroflexi*) showed significant variations among the 4 sites (Fig. 2B). For example, within *Proteobacteria* (Fig 2B, I), *Deltaproteobacteria* abundance increased from 11% to 28% (F_3,30_ = 20.699; p < 0.001) while *Gammaproteobacteria* decreased (19% to 14.8%) from control to 7-year sites (F_3,30_ = 8.803; p < 0.001). Within *Bacteroidetes* (Fig 2B, II), both *Cytophagia* (4% to 8%) and *Bacteroidia* (0.7% to 2.6%) had higher abundances at the 7-year site compared to control sites. The most dramatic increase in abundances was seen in *Firmicutes* (Fig 2B, III) where both *Bacilli* (0.4 to 1.8%) (F_3,30_ = 10.136; p < 0.001) and *Clostridia* (0.9 to 3.3%) (F_3,30_ = 15.418; p < 0.001) increased significantly with years of exposure to oysters. *Chloroflexi* (Fig 2B, IV) showed little variation between control and farmed sites, with only *Chloroflexia* (0.07 to 0.13%) (F_3,30_ = 4.719; p = 0.012) showing a significant increase in 7-year farms sites. Among the top 100 most abundant genera (Appendix S4), several were extremely low in control sites but increased rapidly with duration of farming in the other 3 sites. For example, *Nitrospina* (0.08 to 0.21%), *Desulfobacter* (0.06 to 0.57%), *Desulfotignum* (0.04 to 0.50%), *Natranaerovirga* (0.05 to 0.49%), and *Desulfopila* (0.07 to 0.39%), *Desulfotalea* (0.18 to 0.78%), *Clostriduim* (0.32 to 1.4%), *Bacillus* (0.25 to 1.4%), *Desulfobacterium* (0.71 to 2.6%), *Spirochaeta* (0.44 to 2.9%), *Desulforhopalus* (0.53 to 2.9%), *Draconibacterium* (0.28 to 1.8%) and *Desulfovibrio* (0.39 to 0.78%) all increased from control to 7-year sites. A few genera also showed increased abundances in control sites compared to farmed sites, such as *Pseudomonas* (2.0 to 3.7%), *Phaeodactylum* (0.87 to 2.8%), *Bacillaria* (0.52 to 1.2%), and *Alkanindiges* (0.31 to.75%). Another interesting pattern was observed in some rarer genera, where both *Ca. Brocadia* (F_3,30_ = 6.430; p = 0.003) and *Ca. Scalindua* (F_3,30_ = 4.237; p = 0.018) showed a nearly 3-fold increase in abundance between control and 7-year sites.

To test for differences in community functional attributes, we used PICRUSt2, followed by *ggpicrust2* to analyze the results. PCA analyses of KEGG pathway abundances showed closer clustering of replicates from the farmed sites compared to control with PCA1 axis explaining 26.6% and PCA2 explaining 9.6% of variation in abundances (Fig. 3A). We found 11 KEGG pathways that were statistically different between control and farmed sites, with 8 showing higher (negative fold change) and 3 showing lower enrichment (positive fold change) in farmed compared to control sites (Fig. 3B; p<0.05). Pathways related to antenna proteins (light harvesting proteins in aerobic photosynthesis) and steroid biosynthesis were enriched in control sites and pathways associated with N-glycan biosynthesis, basal transcription factor and mTOR signaling were among the top enriched in farm sites. Using 16S rRNA data to predict community function is limited as putative pathways can be predicted merely due to bacteria containing distant homologs of enzymes important in that pathway, although the pathway itself is not-existent in bacteria (51). Therefore, observations from community RNA analyses are essential to further validate these functional changes.

**Figure 3:**
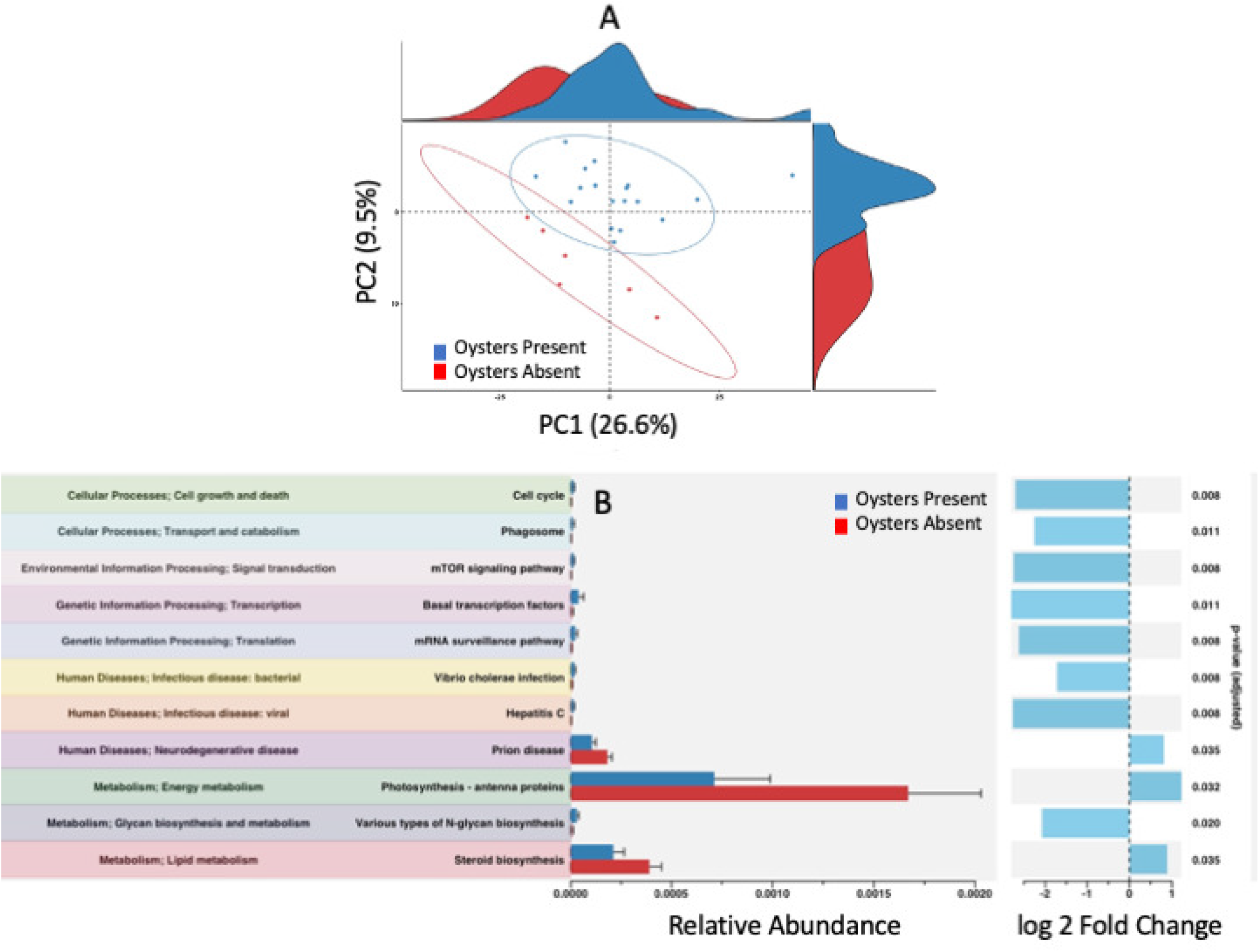
Functional diversity of the bacterial community in control (Oysters Absent: Red) and oyster farms (Oysters Absent: Blue): (A) Principal component analyses and (B) Relative Abundances and log_2_FC of differentially enriched KEGG annotated pathways.

### Sediment Geochemistry

20 elements, excluding hydrogen (H) and oxygen (O), were detected in all 4 sites (Fig. 4; Table 3). nMDS ordination on z-transformed concentrations showed closer clustering between control and 5-year farmed sites, but the former grouped separately from both 3-and 7-year sites (Fig. 4A). PERMANOVA and ANOSIM test indicated a significant difference between sites (Table 2A & B). To identify elements that accounted for these site-specific differences, we performed univariate ANOVAs for individual elements across 4 sites (Fig. 4B). 10 out of 20 elements (Na, K, Cr, Mn, Fe, Co, Ni, Zn, Ba and Pb) showed significant differences in concentrations between sites, where all elements except Na were lower in 7-year site compared to control. Approximately half of the elements (9 out of 20 elements (Na, K, P, Cr, Mn, Fe, Ni, Ba and Pb)) showed significant decreases in concentration in 3-year site compared to control. No elements showed differences in concentrations between control and 5-year site and between 5-year and 7-year site. 3-and 5-year site varied in 5 out of 20 elements (Mn, Fe, Ni, Pb and Cr).

**Figure 4:**
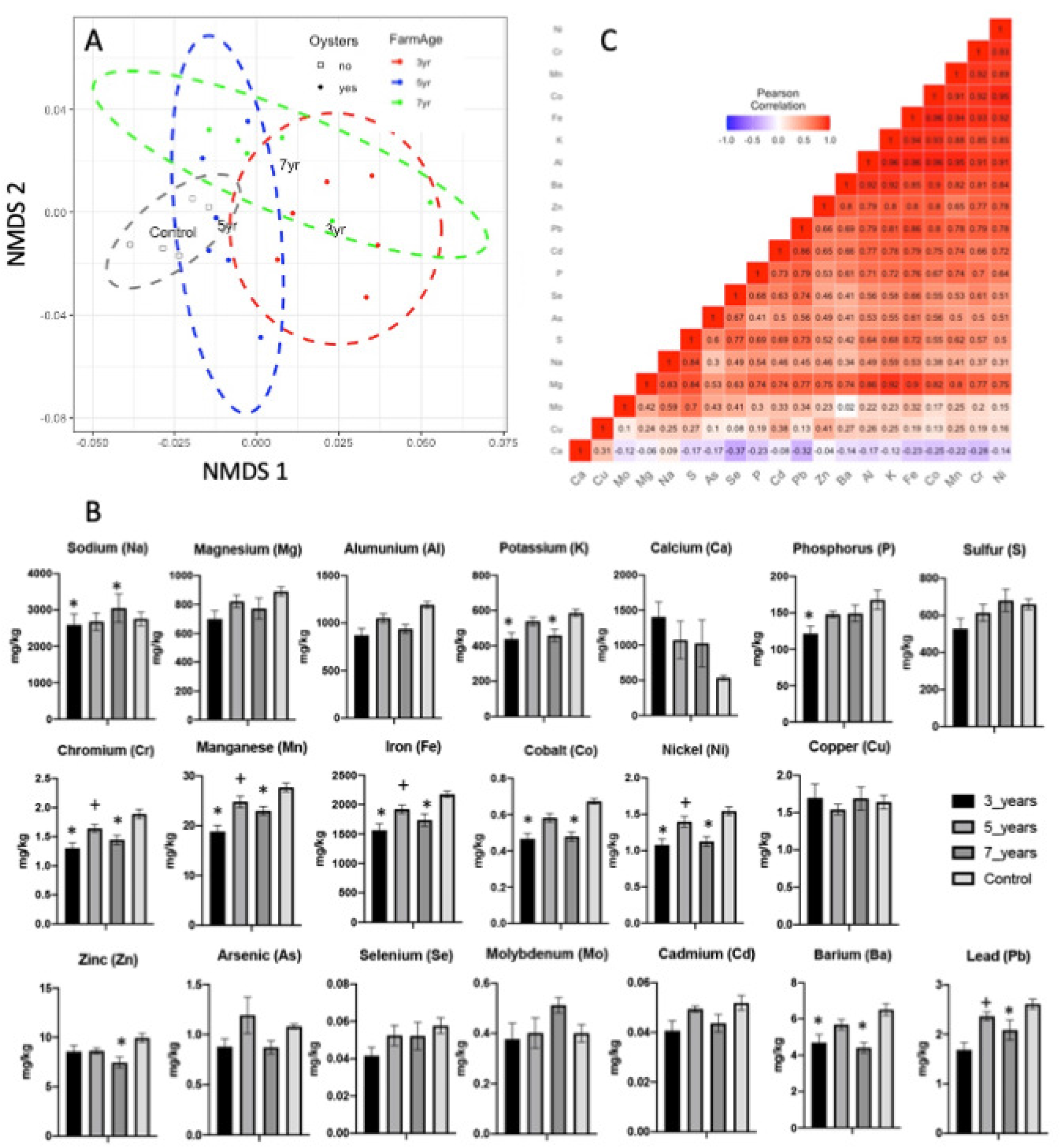
(A) Non-metric multidimensional scaling (nMDS) of elemental concentrations across control, 3-, 5-, and 7-year sites; (B) Concentrations of individual elements across all 4 sampling sites (n= 6). (*) represents significant difference in elements between the respective farm and control sites and (+) represents significant difference in elements between the 3-and 5-year sites (P< 0.05); (C) Pearson Correlation Analyses of elements across all sites

**Table 3:** Concentrations of individual elements in mg/kg across all 4 sampling sites. Data is average±SD, n= 6.

|  | **Metal Concentrations (mg/kg)** |  |  |  |
| --- | --- | --- | --- | --- |
|  | **OA 1 (3 years)** | **OA 2 (5 years)** | **OA 3 (7 years)** | **Control (0 years)** |
| **Sodium (Na)** | 2593.22 ± 734.14 | 2678.81 ± 582.05 | 3051.39 ± 960.92 | 2754.59 ± 469.13 |
| **Magnesium (Mg)** | 699.31 ± 142.77 | 821.64 ± 107.10 | 772.34 ± 179.62 | 890.81 ± 75.63 |
| **Aluminum (Al)** | 873.45 ± 173.79 | 1052.07 ± 117.28 | 939.99 ± 112.36 | 1192.35 ± 87.07 |
| **Potassium (K)** | 438.67 ± 90.27 | 538.70 ± 58.95 | 459.31 ± 87.60 | 586.74 ± 49.52 |
| **Calcium (Ca)** | 1404.57 ± 518.27 | 1074.70 ± 653.45 | 1025.12 ± 818.22 | 536.67 ± 89.48 |
| **Phosphorus (P)** | 121.77 ± 24.24 | 148.07 ± 10.74 | 149.18 ± 27.81 | 167.98 ± 32.51 |
| **Sulphur (S)** | 528.74 ± 134.41 | 614.50 ± 110.69 | 681.09 ± 147.96 | 660.26 ± 70.50 |
| **Chromium (Cr)** | 1.30 ± 0.22 | 1.64 ± 0.16 | 1.44 ± 0.19 | 1.89 ± 0.19 |
| **Manganese (Mn)** | 18.83 ± 2.98 | 24.81 ± 2.82 | 23.01 ± 2.07 | 27.71 ± 2.15 |
| **Iron (Fe)** | 1564.83 ± 271.10 | 1920.93 ± 175.69 | 1737.11 ± 259.50 | 2173.31 ± 137.59 |
| **Cobalt (Co)** | 0.46 ± 0.08 | 0.58 ± 0.058 | 0.48 ± 0.06 | 0.67 ± 0.03 |
| **Nickel (Ni)** | 1.08 ± 0.20 | 1.40 ± 0.19 | 1.12 ± 0.16 | 1.54 ± 0.14 |
| **Copper (Cu)** | 1.69 ± 0.46 | 1.54 ± 0.17 | 1.69 ± 0.38 | 1.63 ± 0.21 |
| **Zinc (Zn)** | 8.59 ± 1.49 | 8.66 ± 0.70 | 7.46 ± 1.36 | 9.94 ± 1.09 |
| **Arsenic (As)** | 0.88 ± 0.19 | 1.19 ± 0.45 | 0.87 ± 0.16 | 1.08 ± 0.06 |
| **Selenium (Se)** | 0.04 ± 0.01 | 0.05 ± 0.01 | 0.05 ± 0.02 | 0.05 ± 0.01 |
| **Molybdenum (Mo)** | 0.38 ± 0.15 | 0.40 ± 0.14 | 0.51 ± 0.076 | 0.40 ± 0.08 |
| **Cadmium (Cd)** | 0.04 ± 0.01 | 0.05 ± 0.00 | 0.044 ± 0.01 | 0.05 ± 0.01 |
| **Barium (Ba)** | 4.70 ± 1.07 | 5.68 ± 0.69 | 4.42 ± 0.70 | 6.51 ± 0.82 |
| **Lead (Pb)** | 1.69 ± 0.36 | 2.36 ± 0.21 | 2.09 ± 0.48 | 2.61 ± 0.24 |

To better understand the relationship between bacterial community and sediment elemental concentrations, Pearson correlation analyses were done to test for collinearity between elements with each other (Fig. 4C) and with bacterial phyla (Fig. 5A). Ca was negatively correlated to all elements except Cu. Ni, Cr, Mn, Co, Fe, K and Al showed the strongest positive correlation with each other compared to the rest (Table X/Figure X). No overall pattern in collinearity was observed between elements and bacterial phyla (Fig. 5A). Pearson correlation coefficient (r) varied from -0.1 to + 0.64, with only *Cyanobacteria* showing a strong positive correlation (>0.3) with K, Co, Ni and Ba. Phyla *Proteobacteria*, *Chloroflexi*, *Fusabacteria*, *Ignavibacteriae*, *Acidobacteria*, *Firmicutes* and *Nitrospinae* showed higher negative r values (>0.3) with correlated elements Cr, Mn, Fe, and Co. K and Ba also showed strong negative correlation with most phyla (Fig 5A). Distance-based redundancy analyses was done to test the contribution of elements on variations of bacterial abundances (Fig. 5B). To check for homogeneity of the bacterial compositional data we first did a detrended correspondence analyses (1^st^ axis length=1.14; 57) followed by PCA with *envfit* function to test for significance of each element. Al, K, Cr, Mn, Fe, Co, Ni, Zn and Ba showed a significant p-value (<0.01) and were used for subsequent RDA analyses (Appendix S5). RDA 1 and RDA 2 together explained 32% of abundance variability (Fig. 5B). Al (p=0.004) and Mn (p=0.025) made a significant contribution on the total variation of bacterial community structure.

**Figure 5:**
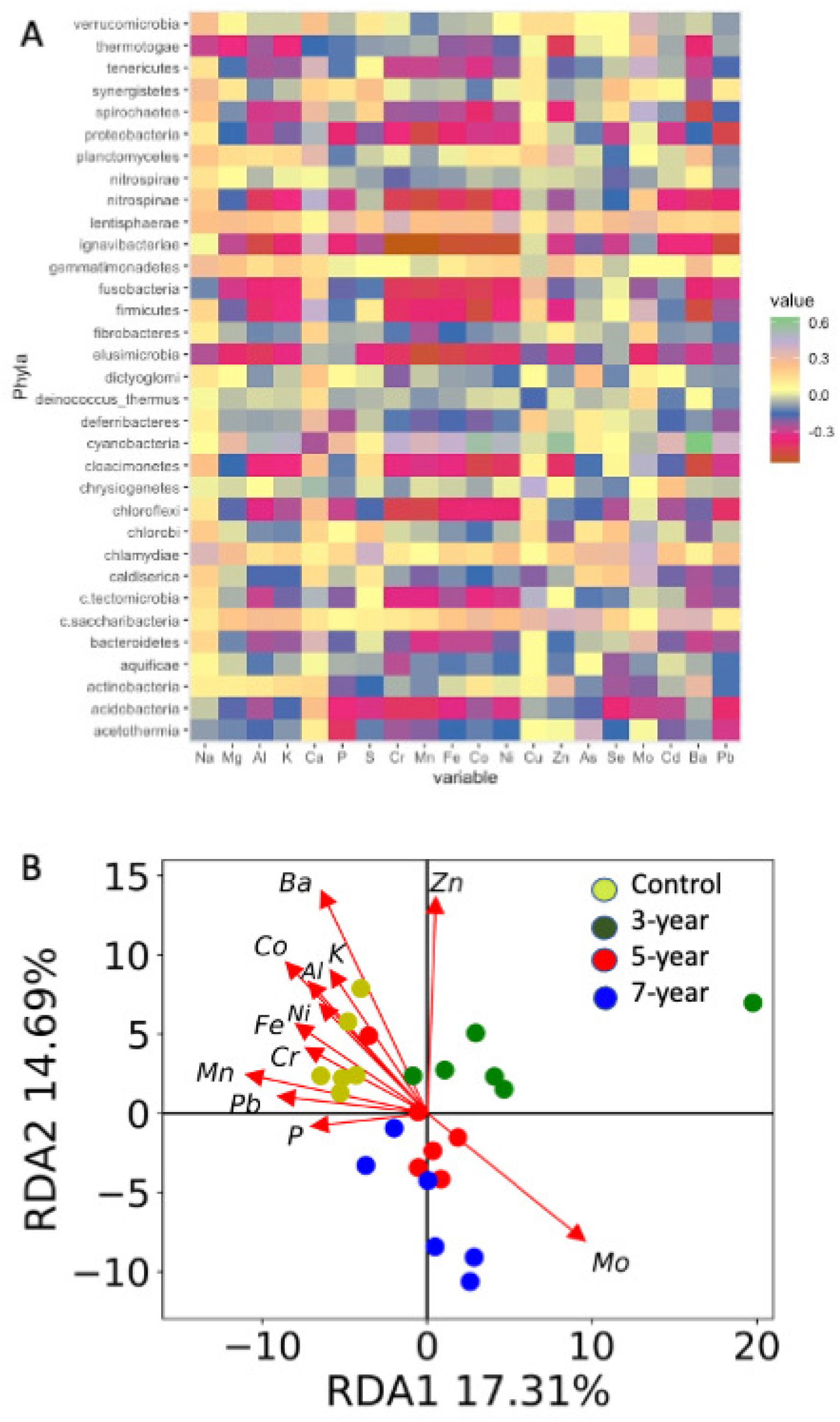
(A) Pearson Correlation Analyses of elements and bacterial phyla across all sites; (B) Biplot of Redundancy Analysis (RDA) axes 1 and 2 for bacterial OTUs with sediment elemental concentrations.

### RNA Community Structure and Function

Illumina HiSeq sequencing of extracted RNA generated an average of 28,084,426 RNA sequences (±12,404,398) in all 6 samples (2 treatments (control vs. 7-year) X 3 replicates; Table 1). No significant difference between RNA yield, RNA quality or sequences were observed between treatment groups (p>0.05). 5459 transcripts were annotated by KEGG that showed no separation in nMDS ordination (Fig. 6A) supported by both PERMANOVA and ANOSIM tests that indicated no differences between control or 7-year site (Table 2A & B). Differential analyses indicated that 8 genes were upregulated, and 18 genes were downregulated in the 7-year site compared to control (Fig. 6B). Among the transcripts upregulated in the control sites, several belonged to gluconeogenesis and bacterial chemotaxis pathway and 1-2 transcripts to phosphotransferase, glutamate, alanine and pyruvate metabolic pathways (Fig. 6C). Multiple pathways related to human diseases, phagosomes and carbon fixation were upregulated in the 7-year farm sites. We annotated our mRNA transcripts with RefSeq in MG-RAST to compare differences in taxonomic assignments from 16S amplicon (resident community) and protein coding reads (active community). We detected a total of 28 and 592 bacterial phyla and genera respectively. nMDS ordination followed by PERMANOVA and ANOSIM on bacterial genera showed no separation between control and 7-year sites (Fig. 7A, Table 2A & B). Diversity analyses on genera also did not show any variation between control and 7-year farm sites (Fig. 7B, Student t-test p>0.01). Similar to 16S amplicons, the top 3 abundant phyla were *Proteobacteria*, F*irmicutes* and *Bacteroidetes*, followed by *Actinobacteria*, *Cyanobacteria*, *Planctomycetes* and *Cloroflexi* (Fig. 7C). 11 out of 28 phyla showed significant increase in farm sites compared to control that included *Proteobacteria*, *Spirochaetes* and *Thermotogae* among the most abundant ones. Only *Planctomycetes* showed a significant increase in control sites. 141 genera showed differential abundance in control and farmed sites (Student t-test; p<0.05), with 124 more abundant in farmed sites and 17 more abundant in control. Supporting 16S amplicon observations, we saw an increase in *Deltaproteobacteria*, *Epsilonproteobacteria* and *Firmicutes* and a decrease in *Alphaproteobacteria*, *<u>G</u>ammaproteobacteria* in the farm sites compared to control. In addition, RNA derived community showed increase of *Actinobacteria, Betaproteobacteria* in the farmed sites and increase of *Planctomycetes* and *Flavobacteria* in the control. To compare similarity of community as predicted by DNA (16S) and RNA derived sequences, we did a Canberra pairwise community distance matrices comparison following protocol described in Meyer et al., (57) (Fig. 7D). For both control and 7-year farm sites, DNA and RNA derived communities were significantly different at the phylum level (Fig. 7D, Control: t=7.64, p<0.001; Farm: t=3.63, p<0.01, Welch Two Sample t-test).

**Figure 6:**
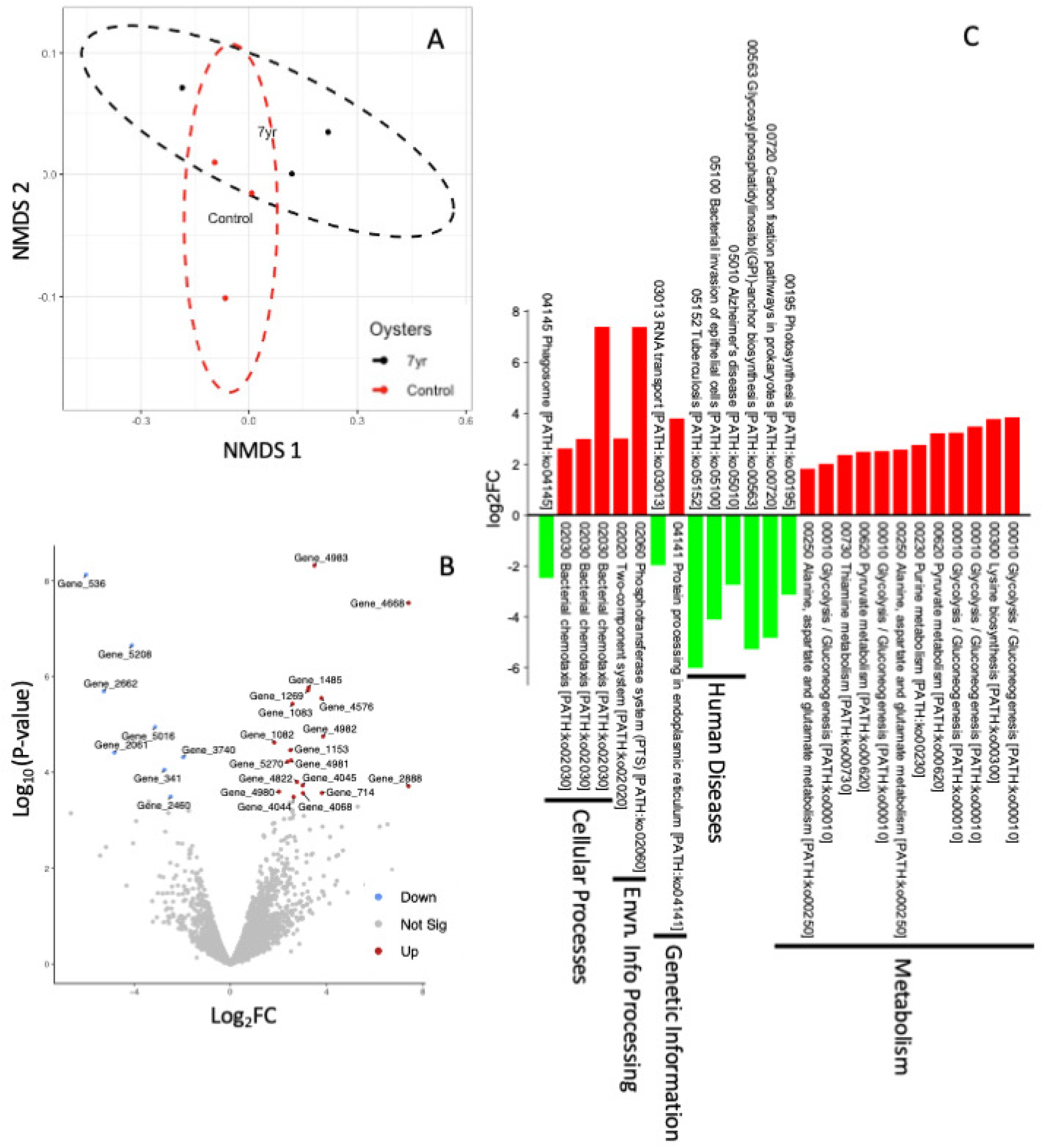
Non-metric multidimensional scaling (nMDS)(A) and Volcano plot of the significant differentially expressed RNA transcripts between control and 7-year sites; (C) Barplot of the differentially expressed KEGG pathways between control and 7-year sites.

**Figure 7:**
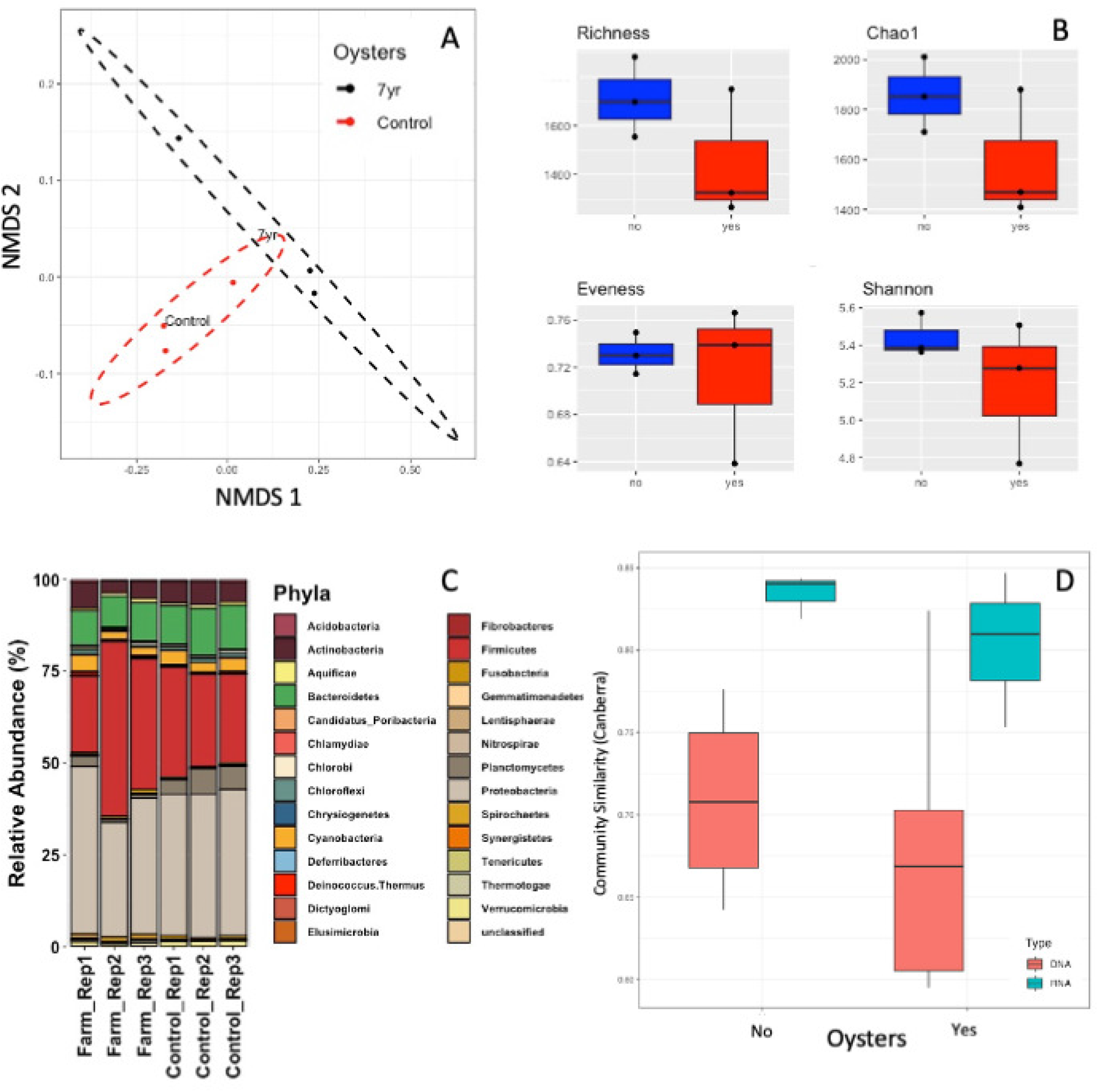
(A) Non-metric multidimensional scaling (nMDS), (B) Diversity Analyses and (C) Relative abundances of bacterial phyla in RefSeq annotated RNA transcripts from control and 7-year sites; (D) Average pairwise similarity (1 -Canberra distance) of the RNA-and 16S amplicon-inferred communities in control (No Oysters) and 7-year (Yes Oysters). Student’s t-test was used to calculate differences between communities (p<0.001).

## Discussion

In this study, we investigated the impact of oyster farming on bacterial community structure and function. We found that bacterial communities in oyster farm sediments differed significantly from unfarmed sediments (Fig. 1 and 2; Table 2). Importantly, we did not find any difference in bacterial communities with respect to farming duration (Fig. 1 and 2; Table 2). In fact, both diversity analyses and linear discriminant analyses (LDA) of effect size (LEfSe) suggest that most bacterial taxa and genera varied in abundances between the control and all three of the farmed sites. These results suggest that sediment bacterial communities responded to the pressure of aquaculture within 3 years, and that this response persists over time. The general lack of change in bacterial communities over farming duration could be due to early and rapid reorganization following the introduction of oyster-induced selection pressures, as observed in studies with similar time frames in marine (58), freshwater (59, 60), and terrestrial systems (61, 62).

Microbial community structure tracks environmental changes due to high adaptation rates (63, 64). As such, microbial communities are increasingly used as indicators to estimate the environmental impacts of anthropogenic activities in different ecosystems (65–67). Aquaculture can impact ecosystem biogeochemistry in several ways, such as decreasing oxygen concentrations (9), increasing sulfide accumulation (22) and enhancing nutrient availability (10). Therefore, we need reliable indicators of these effects to help predict their environmental impacts on coastal systems. In this study, we identified several bacterial taxa that had consistently higher abundance in farmed sites (Fig. 2 and 7). Notable among them is genus *Spirochaeta*, primarily composed of anaerobes found within sediments exposed to aquaculture (68, 69) and class *Clostridia* which are anaerobic species commonly found in soils (70). Additionally, LEfSe (Fig. 2) analyses showed increased LDA scores for sulfate reducing anaerobic *Deltaproteobacteria*, chemolithotrophes associated with *Epsilonproteobacteria* and facultative anaerobes *Bacillales* in the farmed sites. Control sites had higher LDA for *Rhodobacterales* (*Alphaproteobacteria*) that are all strictly aerobic (Fig. 2). RNA derived sequences also showed increase of the anaerobic *Thermotogae* phylum in farmed sites. Ray et al. (36) reported an initial stimulation of sediment O_2_ consumption following the introduction of oyster aquaculture, followed by a return to baseline conditions after several years. Our observed increase in abundance of microbes that prefer, or require anaerobic conditions coupled with past evidence for reduced oxygen availability in sediments under oyster farms suggest a shift towards increased tolerances for anoxia in the oyster farms.

Previous studies show changes in abundance of S cycling bacteria beneath aquaculture (21, 72) due to accumulation of H_2_S in farmed sites (72). LEfSe indicates an overall enrichment of bacterial classes that play significant roles in S cycling, such as *Epsilonproteobactera* and *Deltaproteobacteria* in the farm sites (Fig. 2). Bacterial orders commonly associated with sulfate reduction (e.g., *Desulfobacterales* and *Desulfuromonadales*) within class *Deltaproteobacteria* also showed greater enrichment in farmed sites. Additionally, several sulfur-oxidizing genera (*Desulfopila, Desulfotalea, and Desulfovibrio*) and sulfur-reducing genera (*Desulfobacter* and *Desulfotignum)* (73) showed increased abundances in farms. There is also evidence that oysters may promote sediment PO_4_^3-^ regeneration, but at our sampling site, this was not the case (36). Except an increase in genus *Bacillus,* that includes multiple species with the ability to solubilize inorganic phosphates (PSB) (Fig. 2 (16S) and 7 (RNA)), we found no other taxa that contributes to the coastal P-cycle in both sites (74–76). Remineralization of sediment phosphate is a complex process and is tightly coupled with the Carbon (C):P ratio of the organic substrate (77). Increase in organic matter deposition by oysters in this site can increase the C:P ratio slowing the remineralization rates in farm sites without any microbial mediation.

Oysters play a crucial role in mediating microbial driven nitrogen sediment cycling in estuarine systems. At this site, Ray et al. (78) demonstrated that sediments switched from net nitrogen fixation (i.e., N_2_ sink) to net denitrification (i.e., net N_2_ source) following the addition of oyster aquaculture. Ray (36) also observed a seasonal impact on nitrogen fixation and release rates in these sites. N_2_ can be produced via canonical denitrification, the microbial conversion of nitrate to N_2_, or through anammox, which couples NH_4_^+^ oxidation with NO_2_^-^ reduction to produce N_2_. Ray et al. (36) could not describe the pathway leading to enhanced N_2_ production as they measured net rates. Our analysis of the microbial community and function can help to provide some mechanisms behind the observed change in net N_2_ exchange between sediments and the water column. Here, we see enrichment of few bacterial genus associated with denitrification (e.g., *Bacilli*) and an increased abundance of several anaerobic ammonium oxidizing bacterium (AAOB) capable of anammox including genera *Ca. Brocadia* and *Ca. Scalindua* in the farmed sites (79–81). We also did not see enrichment of bacteria commonly associated with the denitrification processes in the oyster farms (e.g. *Alphaproteobacteria*, *Gammaproteobacteria and Chloroflexi*) (82). *Nitrospinae*, is the only NO_2_^-^ oxidizing bacterial phylum that increased consistently in all farmed sites. Taken together, these data suggest that anammox may have been dominating the N_2_ signal at the farmed sites, potentially because rates of nitrification were lower due to reduced O_2_ availability. Arfken et al. (83) quantified differences in bacterial communities across the digestive system, shell, and sediment in an oyster reef and found that genes associated with denitrification were mostly found in the oyster digestive system and on the shell as opposed to in the sediments further supporting the potential for annamox to dominate N-removal in oyster impacted sediments. We also observed enrichment of phyla *Proteobacteria*, *Firmicutes* and *Spirochaetes*, in farmed sites. All these phyla include taxa with ability to fix nitrogen (84, 85). Fulweiler (86) indicated that N_2_ fixation by can happen for several reasons including (but not limited to) maintaining redox balance in cells and increased mineralization of recalcitrant carbon. Control sites showed enrichment of *Alphaproteobacteria and Gammaproteobacteria*, both, associated with hydrogenotrophic denitrifiers. RNA sequences also showed increased abundance of *Planktomycetes* that has recently been associated with both N_2_ fixing (84) as well as anaerobic ammonium oxidation, or anammox (87). Although the relative contributions of these taxa in driving N-cycle transformations requires further investigations, our data draws attention to the presence of potentially different microbial taxa and processes dominating N-cycle in control versus farmed sites.

Overall, bacterial community composition varied between DNA (resident) and RNA (active) derived sequences (Fig. 7D). Both provided similar estimates of the abundant phyla, except *Actinobacteria* that were overrepresented in the RNA sequences (Fig. 2B & &C), possibly suggesting their increased metabolic activity relative to other groups. Analyzing the RNA derived community help us detect changes in the active community separate from changes in community abundances. For example, RNA sequences showed an increased abundance of *Planctomycetes* in the control sites, that contains many species capable of anaerobic ammonium oxidation, or anammox (87). 16S amplicons showed no such enrichment indicating that although the abundances did not vary between the sites, greater transcriptional efficiency of this group can lead to an active anammox pathway for N_2_ production in the control sites. Several taxa showed similar trends in both 16S and RNA derived community (e.g., *Proteobacteria, Spirochaetes, Clostridia)*, but we also observed taxa that showed no differential abundances in the former but significant differences between sites in the later (e.g., *Flavobacteria*, *Betaproteobacteria*) suggesting that differences in transcriptional rates can contribute to changes in community structure that cannot be predicted from DNA sequences alone. However, compared to 16S taxonomic assignments, RNA transcripts have very limited capacities to predict distribution of rarer taxa in the sediments. Although both sequence data is biased towards certain groups that are overrepresented in the respective databases, our data indicates that bacterial community indices/biomarkers can be effectively used to predict the impact of oyster farming on estuarine systems as proposed in previous studies (21, 71).

Our second objective was to identify the functional status of the microbial community as changes in functional gene transcripts are good indicators of changes in biogeochemical functions (88). Shifts in community structure as seen in the present study (Fig. 1) likely affect the functional capacities of sediments, which in turn can bear significant ecological consequences. Both 16S and RNA transcripts were annotated with the same KEGG database, but no similarity in annotations were observed between the two. No overarching patterns were observed between both DNA and RNA inferred bacterial functions between our control and 7-year sites (Fig. 3, 6; Table 2). Given the inherent variability and the fast turnover rates of mRNA, this is probably not surprising (89), though when looking at expression of individual genes, several were differentially expressed between control and farm sites. We saw a relative increase in abundances of transcripts for antenna proteins that mediates aerobic photosynthesis and transcripts for oxygen-dependent steroid biosynthesis (90) supporting the presence of aerobic conditions in control sites compared to farmed. Enrichment of glutamate (containing assimilated ammonia) and alanine pathways in control sites indicates increased assimilation of nitrogen (91) that is also supported by the increase of *Gammaproteobacteria* and *Planctomycetes* in these sites. Several transcripts involved in glucose production, phosphate metabolism, and bacterial chemotaxis were also enriched in the control sites. The increased organic load and increased inorganic nitrogen availability (36, 78) in the oyster farms probably led to the enrichment of several carbon biosynthesis pathways such as N-glycan biosynthesis and carbon fixation. Several human related pathways (e.g., human disease, phagosomes, basal transcription factors, mTOR signaling) were also enriched, probably indicating a higher degree of anthropogenic disturbances in these farms. For both 16S and RNA data, we were not able to detect changes in genes directly linked to nitrogen, sulphur, and/or anaerobic respiration processes. Additional testing with multiple sites and adequate sequencing depth is necessary to identify reliable functional indicators of change in oyster farms.

Oyster aquaculture can impact elemental concentrations of several metals in sediments due to the correlated nature of most biogeochemical cycles (Fig. 4C; 38, 39). Some studies have indicated an increased concentration of heavy metals (e.g., Cd, Cu, Pb, Zn) in the sediments beneath aquaculture gear due to addition of farm food and increased fecal deposits (23–25), while others have not reported similar effects (38, 92, 93). We predicted higher metal concentrations in farm sites, but instead observed a decrease in most elements from control to 7-year sites (Fig. 4; Table 3). The difference between sites was mostly driven by higher concentrations of elements K, Cr, Mn, Fe, Co, and Ni in the control sites which also exhibited a strong positive correlation with each other. Although further testing is needed to differentiate between natural and aquaculture-derived differences in sediment concentrations, our data indicate that the effect of oyster farms on increased accumulation of metals is negligible in at least in this coastal system. A possible explanation for lower concentrations of several metals beneath aquaculture is a release of metals to the water column under low O_2_ conditions and lower pH in the sediment (94). Further, increased bioaccumulation of metals in oysters could account for reduction in farm sediments, although we have no evidence of that here as we did not measure metal concentrations in oyster biomass (95, 96). Elements Al, K, Cr, Mn, Fe, Co, Ni, Zn and Ba together explained 32% of the bacterial community variations across sites with Al and Mn showing the highest impact (Fig. 5). To our knowledge, our study is the first to show selection effects of metals (Al and Mn) on estuarine bacterial composition, supporting similar observations in river systems (21, 53). Anthropogenic pollution can increase influx of majority of these elements including Al and Mn into estuarine systems (97, 98) which in turn can significantly modify the sediment bacterial communities bearing potential ecological consequences.

Our study used a multi-omics approach to identify changes in the structure and function of sediment microbial communities and associated changes in sediment elemental concentrations across a chrono-sequence (space-for-time substitution) in a commercial oyster farm in Ninigret pond, RI. Our results indicate significant changes in both the structure and function of bacterial communities between the control and farm sites, but this change is not affected by the duration of farming at least up to 7 years as indicated here. We also showed that changes in prominent bacterial taxa can be used as reliable indicators of the impact of oyster farming on estuarine habitats. Differences in bacterial functions between control and farm sites were less pronounced, necessitating further testing and greater sequencing depths to ensure detection of rare but critical functional processes susceptible to farming. We found no evidence of heavy metal accumulation under the oyster cages indicating a trivial effect of oyster farming on estuarine sediment metal concentrations. Such comprehensive analyses of short-and long-term impacts of oyster farming is both novel and crucial to determine the ecological impacts of these practices within the estuarine habitats.

## Acknowledgement

This research was funded, in part, by a grant to R.W.F. from RI Sea Grant and a grant to P.R.C. from NH-INBRE (NIH grant number 5P20GM103506-09, subaward agreement number R1040). R.W.F. and P.R.C. developed the research question. R.W.F. provided funding for sample collection. R.W.F. and P.R.C provided funding for sequencing and laboratory analyses. J.T.E.S. prepared samples for sequencing. We thank Joseph Owens from Keene State College for sample preparation and Tim Maguire and Gabrielle Hillyer from Boston University for assistance with field sampling. NER and ANA also collected samples in the field and performed laboratory analyses of sediment properties. J.T.E.S. and P.R.C. analyzed the data and wrote the paper. All authors edited and revised the paper.

## Data Availability

16S rRNA amplicon DNA and RNA sequences are uploaded to the DDBJ Sequence Read Archive (SRA) under the accession number <u>PRJNA561593</u>.

